# The genomic landscapes of individual melanocytes from human skin

**DOI:** 10.1101/2020.03.01.971820

**Authors:** Jessica Tang, Eleanor Fewings, Darwin Chang, Hanlin Zeng, Shanshan Liu, Aparna Jorapur, Rachel L. Belote, Andrew S. McNeal, Iwei Yeh, Sarah T. Arron, Robert L. Judson-Torres, Boris C. Bastian, A. Hunter Shain

## Abstract

Every cell in the human body has a unique set of somatic mutations, yet it remains difficult to comprehensively genotype an individual cell. Here, we developed solutions to overcome this obstacle in the context of normal human skin, thus offering the first glimpse into the genomic landscapes of individual melanocytes from human skin. We comprehensively genotyped 133 melanocytes from 19 sites across 6 donors. As expected, sun-shielded melanocytes had fewer mutations than sun-exposed melanocytes. However, within sun-exposed sites, melanocytes on chronically sun-exposed skin (e.g. the face) displayed a lower mutation burden than melanocytes on intermittently sun-exposed skin (e.g. the back). Melanocytes located adjacent to a skin cancer had higher mutation burdens than melanocytes from donors without skin cancer, implying that the mutation burden of normal skin can be harnessed to measure cumulative sun damage and skin cancer risk. Moreover, melanocytes from healthy skin commonly harbor pathogenic mutations, likely explaining the origins of the melanomas that arise in the absence of a pre-existing nevus. Phylogenetic analyses identified groups of related melanocytes, suggesting that melanocytes spread throughout skin as fields of clonally related cells, invisible to the naked eye. Overall, our study offers an unprecedented view into the genomic landscapes of individual melanocytes, revealing key insights into the causes and origins of melanoma.

## Introduction

Cutaneous melanomas are skin cancers that arise from melanocytes, the pigment producing cells in the skin. Thousands of melanomas have been sequenced to date, revealing a high burden of somatic mutations with patterns implicating sunlight as the major mutagen responsible for their formation. It is currently unknown precisely when these mutations are acquired during the course of tumorigenesis and whether their rate of accumulation accelerates during neoplastic transformation.

In normal skin, melanocytes reside within the penetrable range of UV-A and UV-B radiation in the basilar epidermis. They make up a minor fraction of the cells in the epidermis, which is mainly comprised of keratinocytes. Keratinocytes have an intrinsic program that triggers apoptosis after exposure to high doses of UV radiation, resulting in the sloughing off of epidermal sheets after a sunburn^1^. This p53-dependent mechanism reduces the persistence of heavily damaged cells in the skin but selects for cells with *TP53* mutations. As a result, clonal patches of *TP53*-mutant keratinocytes are prevalent in sun-exposed skin^2,3^, and these can eventually give rise to keratinocyte cancers.

By contrast, the homeostatic mechanisms governing melanocytes and selective pressures operating on melanocytes during early phases of transformation are less well understood. While some melanomas arise from nevi (i.e. common moles), most arise in the absence of a precursor lesion. Understanding the mutational processes and kinetics of mutation acquisition in pre-malignant melanocytes of normal skin would provide important insights into the early phases of transformation, before clinically visible neoplastic proliferations have formed.

Most DNA sequencing studies are performed on a bulk group of cells, yielding an average signal from the complex mixture of cells that are sampled. Bulk-cell sequencing of normal blood^4^, skin^3^, esophageal mucosa^5^, and colonic crypts^6^ has identified mutations in these tissues, including the presence of pathogenic mutations, offering valuable insights into the earliest phases of carcinogenesis in these tissue types. However, bulk-cell sequencing is not suited to detect mutations in melanocytes because melanocytes are sparsely distributed in the skin^3^.

Genotyping studies at the resolution of individual cells are rare and none have been performed on melanocytes. Genotyping an individual cell is difficult because there is only one molecule of dsDNA corresponding to each parental allele in a diploid cell. There are primarily two strategies to overcome this bottleneck. First, an individual cell can be sequenced after amplifying its genomic DNA *in vitro*^7,8^. Unfortunately, *in vitro* amplification regularly fails over large stretches of the genome, reducing the sensitivity of mutation detection, and errors are frequently incorporated during amplification, diminishing the specificity of subsequent mutation calls^9^. Alternatively, a cell can be clonally expanded in tissue culture, prior to sequencing, to increase genomic starting material^10–13^, but only limited types of primary human cells can sufficiently expand in tissue culture, reducing the scope of this strategy. Here, we combine elements of each strategy, allowing us to genotype melanocytes from normal skin at single-cell resolution.

## Results

### A workflow to genotype individual skin cells

We collected clinically normal skin from 19 sites across 6 donors. Skin biopsies were obtained from cadavers with no history of skin cancer or from peritumoral tissue of donors with skin cancer (Fig. 1A). All donors were of light skin tone, European ancestry (Fig S1A), and ranged from 63 to 85 years in age.

**Figure 1.**
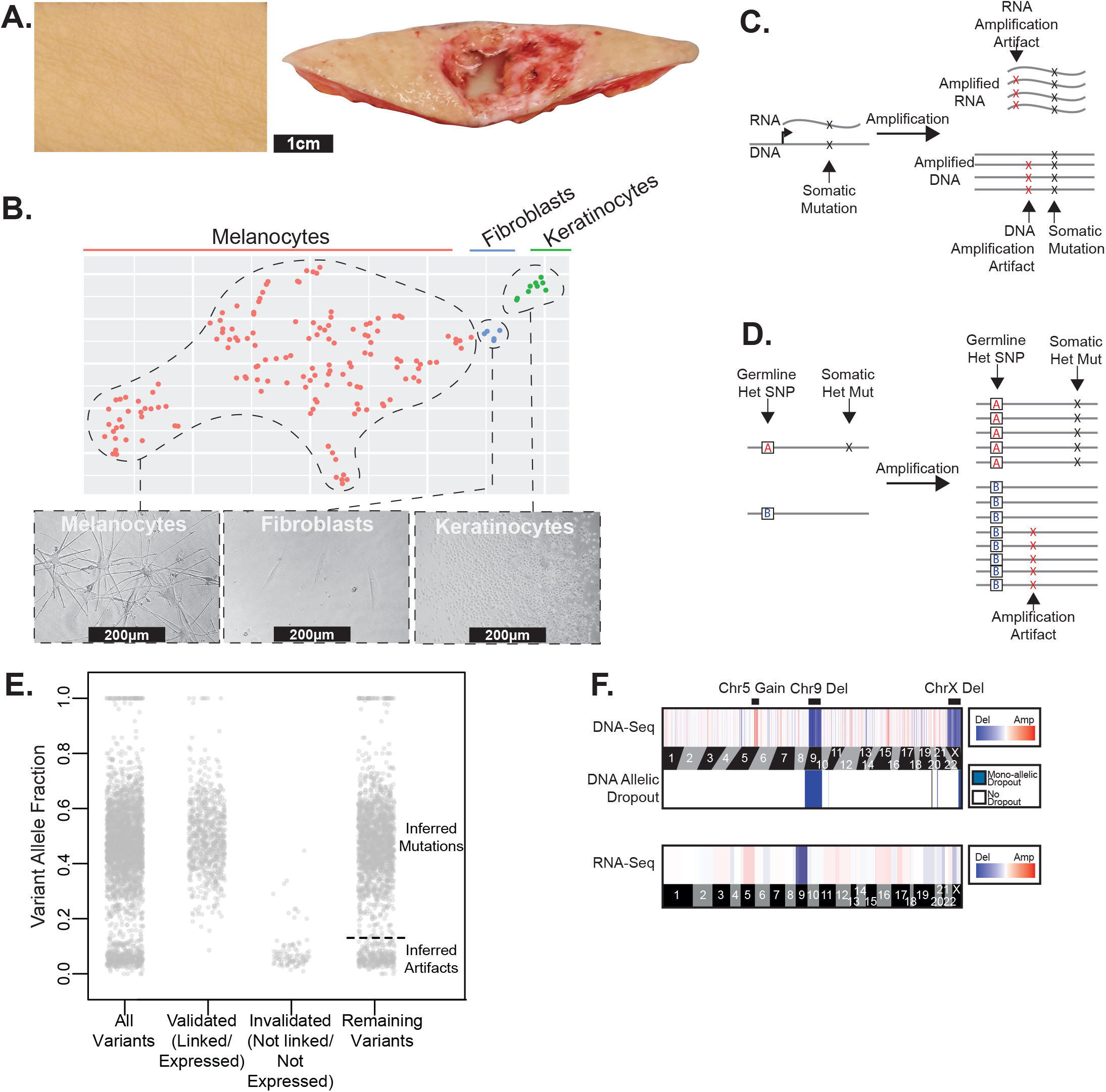
A workflow to genotype individual skin cells. **A.** Examples of healthy skin from which we genotyped individual cells. Left panel: skin from the back of a cadaver. Right panel: skin surrounding a basal cell carcinoma. **B.** Expression profiles classify the cells that we genotyped into their respective lineages. Cells are depicted in a t-SNE plot and colored by their morphology. See Fig. S1B for further details. **C-D.** Patterns to distinguish true mutations from amplification artifacts. **C.** Mutations in expressed genes are evident in both DNA- and RNA-sequencing data, whereas amplification artifacts are not. **D.** Germline polymorphisms, distinguished here as “A” and “B” alleles, are in linkage with somatic mutations but not amplification artifacts. **E.** Variant allele fractions from an example cell indicate how we inferred the mutational status of variants outside of the expressed and phase-able portions of the genome. Variants that were validated as mutations had variant allele fractions (VAFs) around 1 or 0.5, and variants that were invalidated had lower VAFs; however, PCR biases sometimes skewed these allele fractions. Variants that could not be directly validated or invalidated were inferred by their VAF (see methods for details). The dotted line indicates the optimal VAF cut-off to distinguish somatic mutations from amplification artifacts for this particular cell’s variants (see Fig. S2B for more details). **F.** Copy number was inferred from DNA- and RNA-sequencing depth as well as from allelic imbalance--an example of a cell with a gain over chr. 5q, loss of chr. 9, and loss of chr. X is shown.

From each skin biopsy, epidermal cells were established in tissue culture for approximately two weeks and subsequently single-cell sorted and clonally expanded. On average, 38% of flow-sorted melanocytes produced colonies, ranging from 2-3000 cells (median 184 cells, Table S1), indicating that we are studying a prevalent and representative population. Despite the small size of these colonies, there was sufficient starting material to eventually achieve 99.86% allelic coverage (i.e. 0.14% allelic dropout) (Fig. S2A).

Next, we extracted, amplified, and sequenced both DNA and RNA from each clonal expansion, as described^14,15^. Our tissue culture conditions were tailored for melanocyte growth, but some keratinocytes and fibroblasts also grew out. The RNA sequencing data provided confirmation of each cell’s identity (Fig. 1B, S1B). The matched DNA/RNA sequencing data also permitted genotype/phenotype inquiries, as described in subsequent sections.

Polymerases often introduce errors during amplification, and these artifacts can be difficult to distinguish from somatic mutations. The matched DNA/RNA sequencing data improved the specificity of mutation calls because mutations in expressed genes could be cross-validated, whereas amplification artifacts arise independently during DNA and RNA amplifications and thus do not overlap (Fig. 1C). To further improve the specificity of mutation calls, we leveraged haplotype information to root out amplification artifacts. When reads are phased into their maternal and paternal haplotypes using heterozygous germline variants, neighboring somatic mutations occur within all amplified copies of that haplotype, whereas amplification artifacts rarely display this pattern (Fig. 1D)^16,17^.

Altogether, we were able to confidently distinguish true somatic mutations from amplification artifacts in portions of the genome that were expressed and/or could be phased. Variants that fell outside of these regions were classified as somatic mutations or artifacts based on their variant allele frequencies. Heterozygous mutations should have allele frequencies of 50%, whereas, amplification artifacts tend to have much lower allele frequencies. For each cell, we identified the variant-allele-frequency cutoff that would maximize the specificity and sensitivity of somatic mutation calls by comparing the variant allele frequencies of the known somatic mutations and the known amplification artifacts in the expressed and phase-able portions of the genome (Fig 1E, S2B-D).

Finally, we deduced copy number alterations from both the DNA-seq data and the RNA-seq data using the CNVkit software suite^18,19^. As an additional filter, we required that copy number alterations coincide with a concordant degree of allelic imbalance over the region affected (Fig. 1F).

In summary, we implemented a series of experimental and bioinformatic solutions to overcome the major obstacles associated with genotyping individual melanocytes. 133 melanocytes passed our quality control metrics and were included in all subsequent analyses. Tissue pictures, cellular morphologies, and genomic features (point mutations, copy number alterations, and allelic imbalance information) are shown for each melanocyte in a supplemental dataset hosted by figshare (https://figshare.com/s/bb5614d5ab4554516278)^20^.

### Mutational landscape of melanocytes from normal skin

For each clone, we performed RNA-sequencing of the entire transcriptome and DNA-sequencing on a panel of 509 cancer-relevant genes (see Table S2 for baits). For a subset of 48 cells with low mutation burdens we performed an additional round of DNA-sequencing over the entire exome, providing more power to measure the mutational signatures operating in those cells.

We observed an average mutation burden of 7.9mut/Mb (mutations per megabase); however, this ranged from 0mut/Mb to 32.3mut/Mb, depending upon several factors (see Table S3 for mutation calls). The mutation burdens of melanocytes first varied within people by anatomic site. As expected, melanocytes from sun-shielded sites had fewer mutations than those on sun-exposed sites (Fig. 2A,B). Consistently, sun-shielded melanocytes had little evidence of UV-radiation-induced mutations, whereas, this was the dominant mutational signature in melanocytes from sun-exposed skin (Fig. 2A).

**Figure 2.**
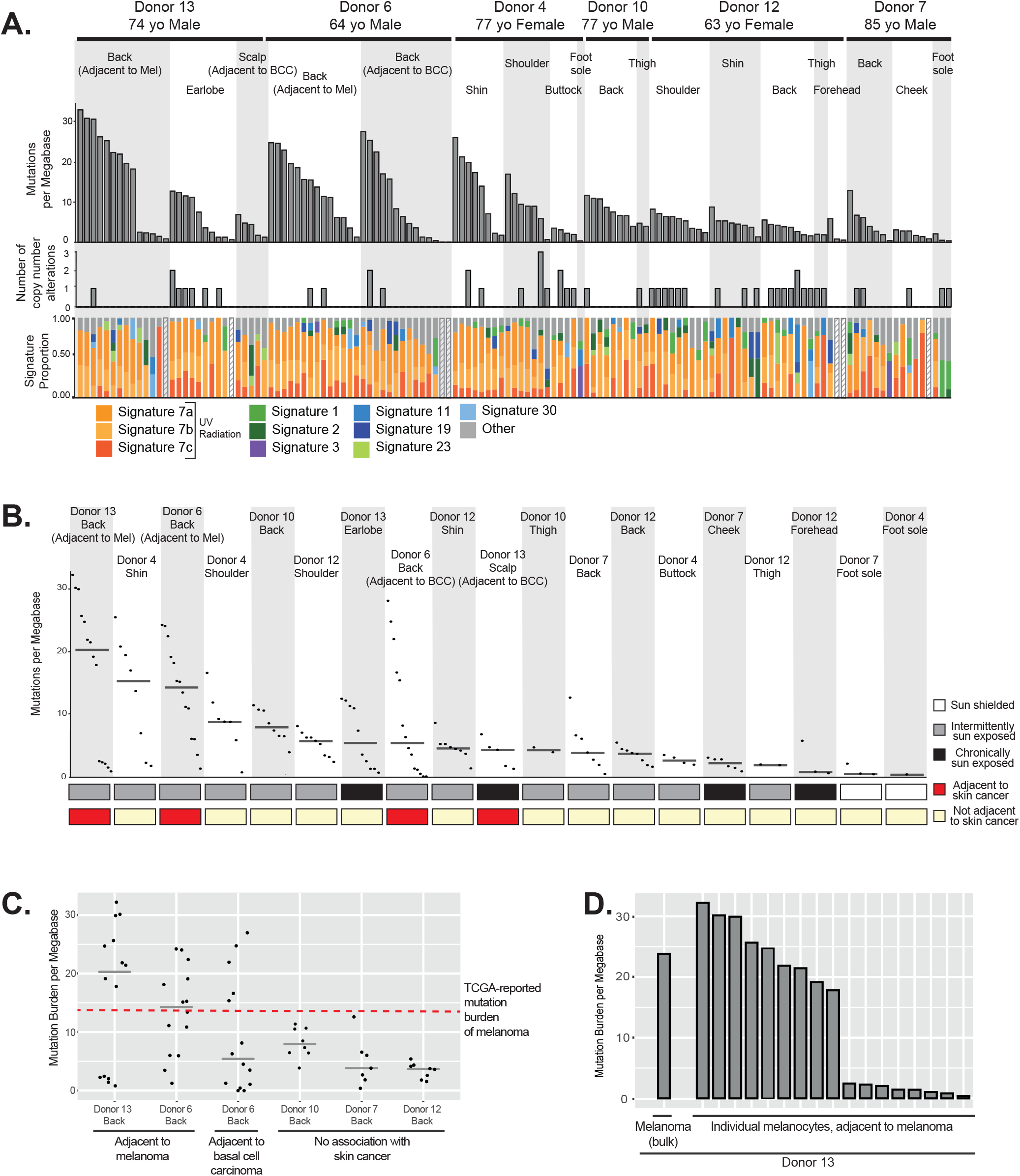
The genomic landscape of individual melanocytes from physiologically normal human skin. **A.** Top panel: Mutation burden of melanocytes from physiologically normal skin of six donors across different anatomic sites (BCC = Basal Cell Carcinoma, Mel = Melanoma). Middle panel: Number of copy number alterations identified within each melanocyte. Lower panel: The proportion of each cell’s mutations that are attributable to established mutational signatures. Signatures 7a, 7b and 7c are associated with UV-radiation induced DNA damage. Hashed bars indicate that there were too few mutations for signature analysis. **B.** Mutation burden plotted again but rank ordered by median mutation burden (line) within each site. **C.** Mutation burden of melanocytes from back skin adjacent to cancer in contrast to back skin not adjacent to cancer. The median is denoted by a grey line, and the red dotted line denotes the mutation burden of melanoma. **D.** Mutation burden of melanocytes as compared to an adjacent melanoma.

Surprisingly, among sun-exposed melanocytes, cells from the back and limbs had more mutations than cells from the face (Fig. 2A,B). Typically, skin from the back and limbs is only exposed intermittently to sunlight and expected to accumulate lower levels of cumulative sun exposure than skin from the face, neck, and bald scalp. The finding of lower mutation burdens in chronically sun-exposed sites deserves further study as it indicates possible differences in mutation rate, DNA repair or turnover among melanocytes from these anatomic sites. However, our observations are consistent with the fact that melanomas are disproportionately common on intermittently sun-exposed skin as compared to other forms of skin cancer^21,22^.

The mutation burdens of melanocytes also varied between donors. For example, we sequenced melanocytes from a common site, the back, of five donors. Among these, the melanocytes from donors 6 and 13 harbored the highest mutation burdens (Fig. 2C) with more than half of melanocytes exceeding the average mutation burden of melanoma (14.4 mut/Mb^23^) – this was notable because these donors had skin cancer adjacent to the skin that we sequenced.

For several donors, we observed a wide range of mutation burdens among the melanocytes harvested from the same anatomic site. This is surprising as cells originating from the same area of skin (~3cm^2^) would be expected to have similar levels of exposure to UV radiation and therefore comparable mutagenic profiles. To further understand the broad range of mutation burden, we sought to identify genes whose expression correlates with mutation burden using differential expression analysis (Fig. S3). Among the significant genes, *MDM2* was more highly expressed in melanocytes with elevated mutation burdens. MDM2 promotes the rapid degradation of p53, raising the possibility that there is heterogeneity among melanocytes with respect to p53 activity, which could affect the ability of a cell to repair mutations or undergo DNA damage-induced cell death. Although MDM2 provides a convincing narrative to explain the mutation burden heterogeneity, it is just one out of a number of significantly correlated genes that may be contributing to the phenotype. Another possibility is that the melanocyte population in our sample is heterogeneous. It could comprise of melanocytes with different residence times in the epidermis or different developmental states. For example, the low mutation burden melanocytes may reside, or have resided for some portion of their life, in a privileged niche, such as the hair follicle, thereby protecting them from UV-radiation. Future studies will be needed to better resolve why melanocytes from a single site can exhibit such a broad range of mutation burdens.

Melanocytes were harvested near a site with melanoma in two patients, and tumor tissue was available from one of the donors. The mutation burden of the melanoma, determined by bulk sequencing, was comparable to that of the individual melanocytes from its surrounding skin (Fig. 2D). There was no overlap between mutations in the melanoma and surrounding melanocytes, ruling out the possibility that tumor cells contaminated the normal melanocyte population. While more cases need to be studied, our findings suggest that melanomas have mutation burdens similar to their neighboring normal cells. This would contrast with colorectal cancers, which have higher mutation burdens than surrounding normal colorectal cells^24^.

Copy number alterations were relatively uncommon in melanocytes (Fig. 2A middle panel, S4), with the exception of recurrent losses of the Y-chromosome and the inactive X-chromosome (Table S4). Mosaic loss of the Y-chromosome and the inactive X-chromosome has been reported in normal blood^25,26^, suggesting that this is a generalized feature of aging. The rarity of autosomal copy number alterations in melanocytes from normal skin is consistent with previous reports that copy number instability is acquired during the later stages of melanoma evolution, and thus unlikely to be operative in pre-neoplastic melanocytes^27,28^.

### Pathogenic mutations in melanocytes from normal skin

We next explored the mutations to identify those that have been previously attributed as drivers of neoplasia. A set of 29 pathogenic mutations were identified in 24 different cells (Fig. 3A). In particular, there were numerous mutations predicted to activate the Mitogen-Activated Protein Kinase (MAPK) pathway. These include loss-of-function mutations in negative regulators of the MAPK pathway, affecting *NF1*, *CBL*, and *RASA2*. There were also gain- or change-of-function mutations in *BRAF*, *NRAS*, and *MAP2K1*, however, *BRAF^V600E^* mutations – the most common mutation in the MAPK pathway occurring in melanocytic neoplasms^23,29^ – were not detected.

**Figure 3.**
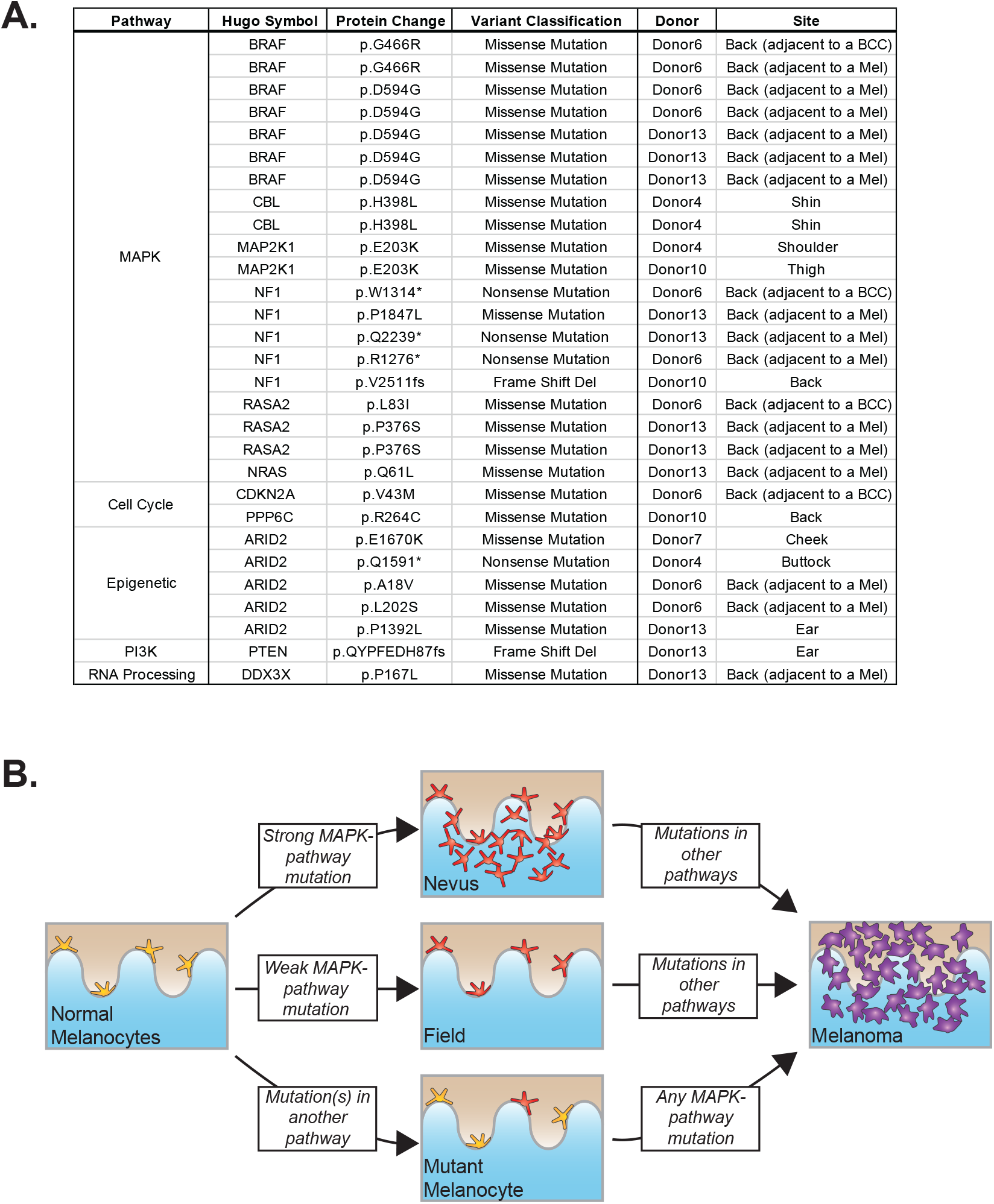
Pathogenic mutations in melanocytes from normal human skin. **A.** A curated list of pathogenic mutations in melanocytes found in this study. BCC = Basal Cell Carcinoma, Mel = Melanoma. **B.** Based on the data shown here and in conjunction with previous genetic, clinical, and histopathologic observations, we propose that melanomas can evolve via distinct trajectories, depending upon the order in which mutations occur. MAPK = Mitogen-Activated Protein-Kinase.

The World Health Organization (WHO) classification of melanoma distinguishes two major subtypes of cutaneous melanoma – the low cumulative sun damage (low CSD) subtype and the high cumulative sun damage (high CSD) subtype. Low CSD melanomas are driven by *BRAF^V600E^* mutations and often originate from nevi^30^. The *BRAF^V600E^* oncoprotein is highly active and likely sufficient to form a melanocytic nevus, explaining the absence of these mutations in skin without evident melanocytic neoplasia. By contrast, high CSD melanomas are driven by a more diverse set of mostly weaker MAPK pathway mutations, and they arise *de novo* rather than from nevi^30^. The pathogenic mutations found in our study overlapped with those known to occur in high CSD melanomas and thus likely stem from incipient neoplastic clones that could eventually progress to high CSD melanomas (Fig. 3B).

We also observed driver mutations in other signaling pathways, including mutations that disrupt chromatin remodeling factors and cell-cycle regulators (Fig. 3A). These mutations are presumably not sufficient to induce a neoplasm but likely accelerate progression in the event that the harboring cell acquires a MAPK pathway mutation^31^ (Fig. 3B). These melanocytes thus represent a poised cell that could immediately transform into a rapidly growing neoplasm, skipping the incremental phases of tumor progression, once a mutation in the MAPK pathway occurs. This evolutionary trajectory may explain the evolution of nodular melanoma, a type of melanoma that occurs in the absence of a nevus and grows rapidly^32^.

### Melanocytes can persist as fields of related cells within the skin

We found shared mutations between nine separate sets of melanocytes, suggesting that these cells stem from clonal fields of melanocytes in the skin (Fig. 4). We ruled out the possibility that these melanocytes emerged during our brief period of tissue culture by growing neonatal melanocytes for several months and measuring their mutation burdens over time (see methods for a full description) (Fig. S5). The number of private mutations in the related sets of melanocytes, shown in figure 4, was many orders of magnitude higher than would be expected from two weeks in tissue culture. Moreover, the private mutations from sun-exposed melanocytes showed evidence of UV-radiation-induced DNA damage (Fig. 4) – a mutational process that does not operate in tissue culture^33^.

**Figure 4.**
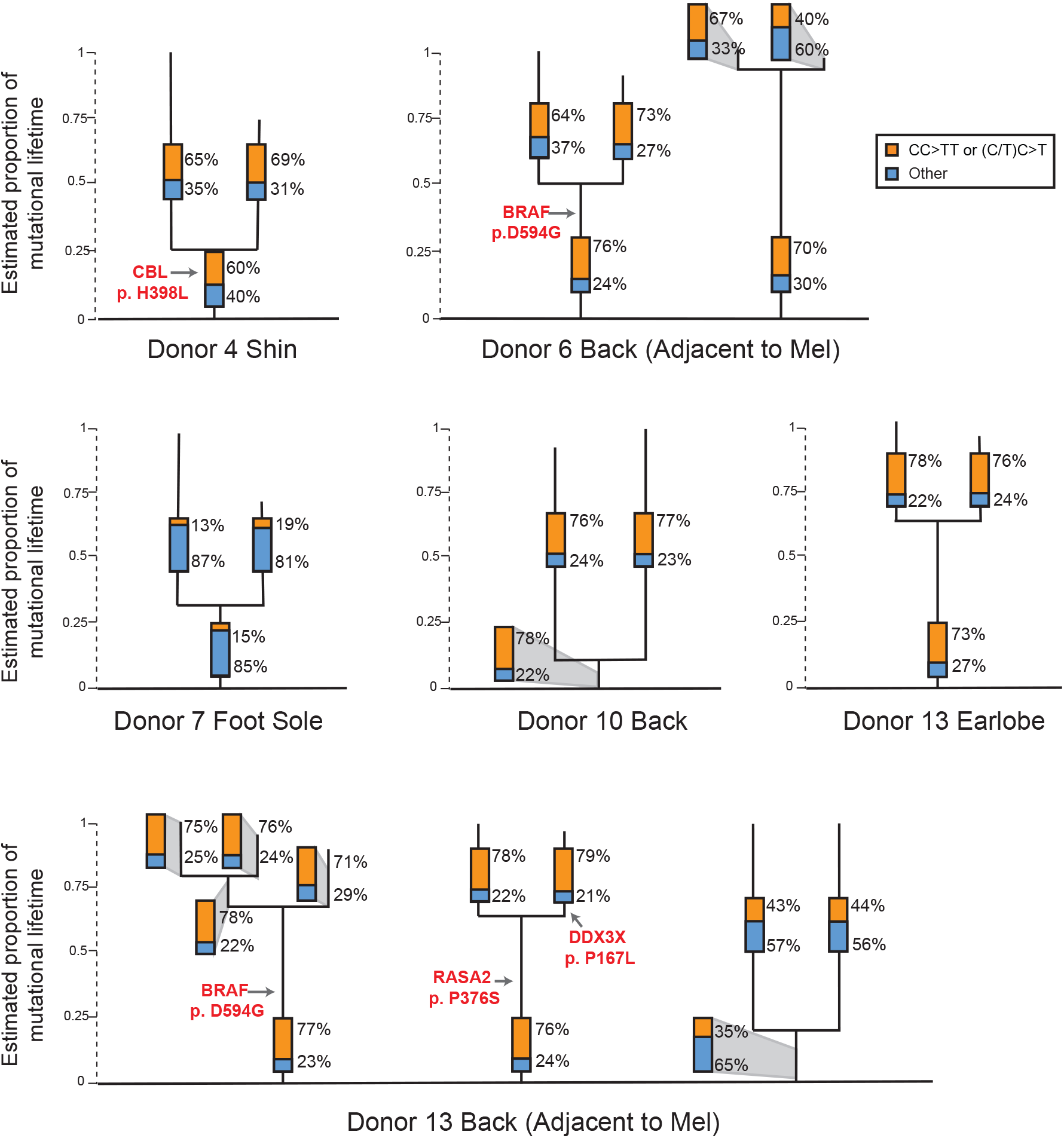
Fields of related melanocytes identified in normal human skin. Phylogenetic trees in which each line terminus corresponds to an individual cell. Mutations that are shared between cells comprise the trunk of each tree and private mutations in each cell form the branches. Trunk and branch lengths are scaled equivalently within each tree but not across trees. The proportion of mutations that can be attributed to ultraviolet radiation (CC>TT or (C/T)C>T) is annotated in the bar charts on each tree trunk or branch. Pathogenic mutations and their location on each tree are indicated in red text. Mel = Melanoma.

Four of the sets of related melanocytes harbored a pathogenic mutation in the trunk of their phylogenetic trees, implicating the mutation in the establishment of the field. It is possible that the remaining fields of melanocytes had a pathogenic mutation that we did not detect or appreciate, but we favor the explanation that fields of related melanocytes can also form naturally over time, for instance, as the body surface expands or as part of homeostasis.

## Discussion

There is a complex set of risk factors associated with melanoma, including cumulative levels of sun exposure, peak doses and timings of exposures throughout life, skin complexion, tanning ability, and DNA repair capacity^34^. It is nearly impossible to quantify and integrate the effects of each of one of these variables, but we demonstrate here that it is feasible to directly measure the mutational damage in individual melanocytes. Moving forward, the number and types of mutations in melanocytes warrant further exploration as biomarkers to measure cumulative sun damage and melanoma risk.

Our study also offers important insights into the origins of melanoma. Idealized progression models typically depict melanomas as passing through a series of precursor stages, but in reality, most melanomas appear suddenly, without an association to a precursor lesion^35^. We show that human skin is peppered with individual melanocytes or fields of related melanocytes harboring pathogenic mutations known to drive melanoma. These poised melanocytes likely give rise to melanomas that appear in the absence of a pre-existing nevus, once additional mutations are acquired.

Finally, our genomic studies are an important resource to further understand basic melanocyte biology. For example, we found that melanocytes from sun-damaged skin vary in their mutation burdens by multiple orders of magnitude. Of note, a similar pattern of variable mutation burdens was recently reported in bronchial epithelial cells of former smokers^13^. Melanocytes with few mutations are likely to be more efficient at DNA repair and/or have occupied privileged niches, protected from the sun, such as in the hair follicle. Melanocyte stem cells in the hair follicle can contribute to the intraepidermal pool of melanocytes as is evident in vitiligo patients with repigmenting areas^36^ – a similar process may be operative in the general population to replenish sun-damaged melanocytes.

In summary, the genetic observations described here offer new insights into the early phases of melanocytic neoplasia, melanocyte homeostasis, and the consequences of UV radiation. Our results suggest that mutation burden in normal melanocytes may serve as biomarkers to measure sun damage, assess the effectiveness of interventions such as sunscreens, and provide refined prediction of an individual’s melanoma risk.

## Acknowledgements

We acknowledge support from the following: National Cancer Institute – K22 CA217997 (AHS), Melanoma Research Alliance (AHS), LEO Foundation (AHS), George and Judy Marcus Precision Medicine Fund (AHS and STA), National Center for Advancing Translational Sciences and the National Institutes of Health through UCSF-CTSI TL1-TR001871 (JT), 1R35CA220481 (BCB), Mt. Zion Health Research Fund (AHS), Dermatology Foundation (AHS), the American Federation of Aging Research (AHS), and the NIH Director’s Common Fund – DP5 OD019787 (RLJ). We thank the tissue donors, whose tissue was obtained through the UCSF Willed Body Program for medical education, and patients who consented to donate surgical discard tissue. Cell sorting was performed in the Laboratory for Cell Analysis of UCSF’s Helen Diller Family Comprehensive Cancer Center which is supported by a National Cancer Institute Cancer Center Support Grant (P30 CA082103).

## Author Contributions

Conception and design of the work: A. Hunter Shain. Data collection: Jessica Tang, Darwin Chang, Shanshan Liu, Eleanor Fewings, Hanlin Zeng, Aparna Jorapur, Rachel L. Belote, Andrew S. McNeal, Sarah T. Arron. Data analysis and interpretation: Eleanor Fewings, Jessica Tang, Darwin Chang, Robert L. Judson-Torres, Boris C. Bastian, A. Hunter Shain. Drafting the article: Eleanor Fewings, Jessica Tang, A. Hunter Shain. Critical revision of the article: Eleanor Fewings, Jessica Tang, A. Hunter Shain, Rachel L. Belote, Iweh Yeh, Sarah T. Arron, Robert L. Judson-Torres, Boris C. Bastian.

---

**Figure S1.**
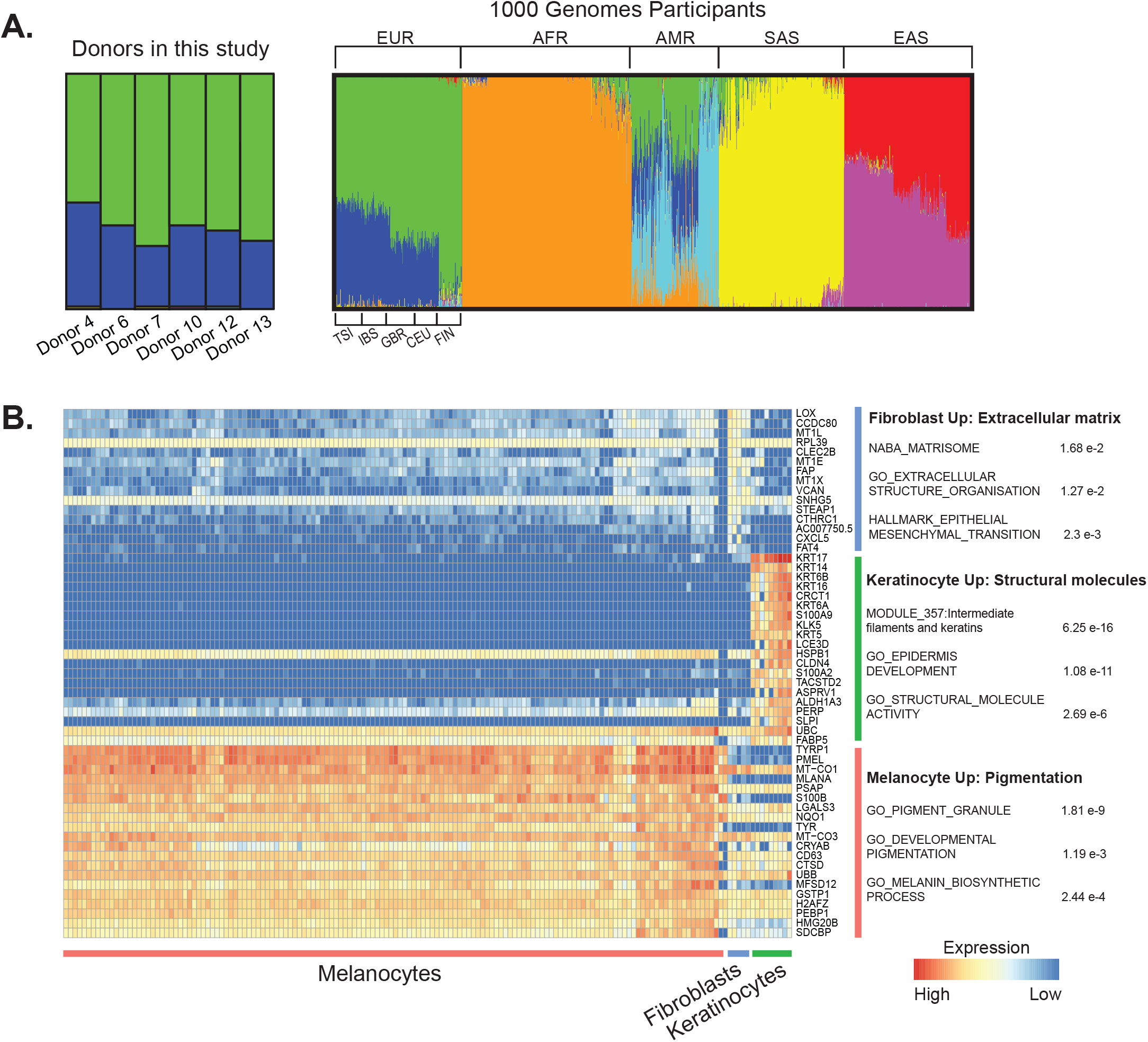
Establishing the ethnicity of donors and identity of cells in this study. **A.** Admixture analysis of donors included in this study alongside participants from the 1000 Genomes Project. Donors in our study were genotypically most similar to European participants from the 1000 Genomes Project. EUR-European (TSI-Toscani in Italia, IBS - Iberian Population in Spain, GBR - British in England and Scotland, CEU - Utah Residents with Northern and Western European Ancestry, FIN - Finnish in Finland), AFR - African, AMR - Latin American, SAS - South Asian, and EAS - East Asian. **B.** Differential expression analysis comparing cells that were morphologically predicted to be keratinocytes, melanocytes, or fibroblasts (see Fig. 1B for more details). The top 20 differentially expressed genes for each group are shown along with gene ontology terms with significant overlap.

**Figure S2.**
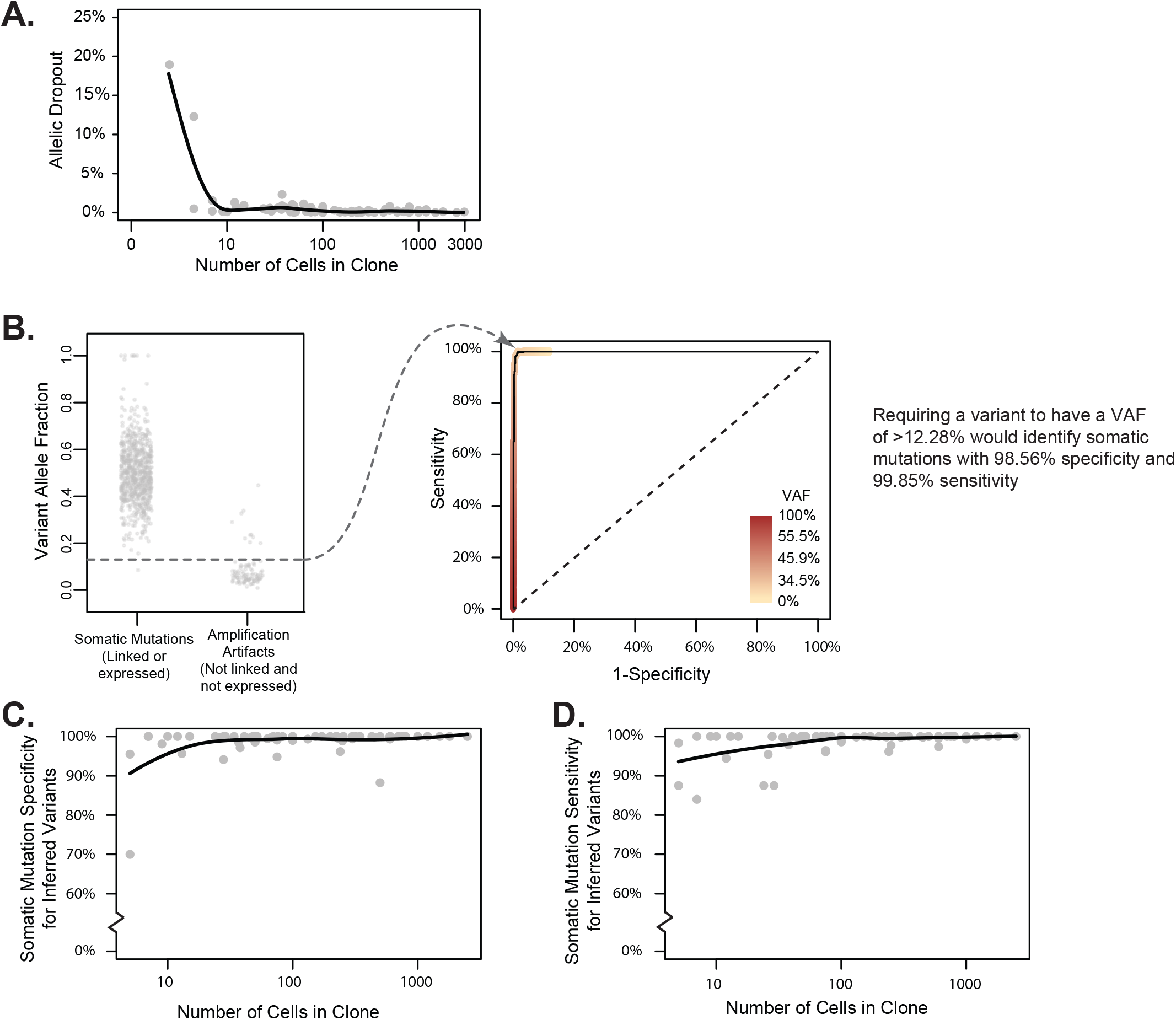
Detection of somatic mutations in small clones of skin cells with high specificity and sensitivity. **A.** Allelic dropout declines rapidly as a function of clone size. Each data point represents the percent of germline SNP alleles that could not be detected for a given clone as a function of the number of cells within the clone. **B.** Establishing a variant allele fraction (VAF) cut-off to infer somatic mutations within a clone. The left panel depicts the VAFs for known somatic mutations and known amplification artifacts from a single clone. The right panel depicts a ROC curve, showing the VAF at which sensitivity and specificity of somatic mutation calls would be maximized when inferring the mutational status of variants based on VAF alone. Variants that fell within expressed or phaseable portions of the genome were classified as mutations or artifacts as described (see Fig. 1C,D). The remaining variants were inferred based on a VAF cut-off, which maximized sensitivity and specificity of somatic mutation calls, as shown here. **C-D.** The specificity (panel C) and sensitivity (panel D) of inferred somatic mutations as a function of clone size. The mean specificity and sensitivity of inferred somatic mutations was respectively 98.83% and 98.60% for all clones of at least 5 cells. All trendlines correspond to a moving average.

**Figure S3.**
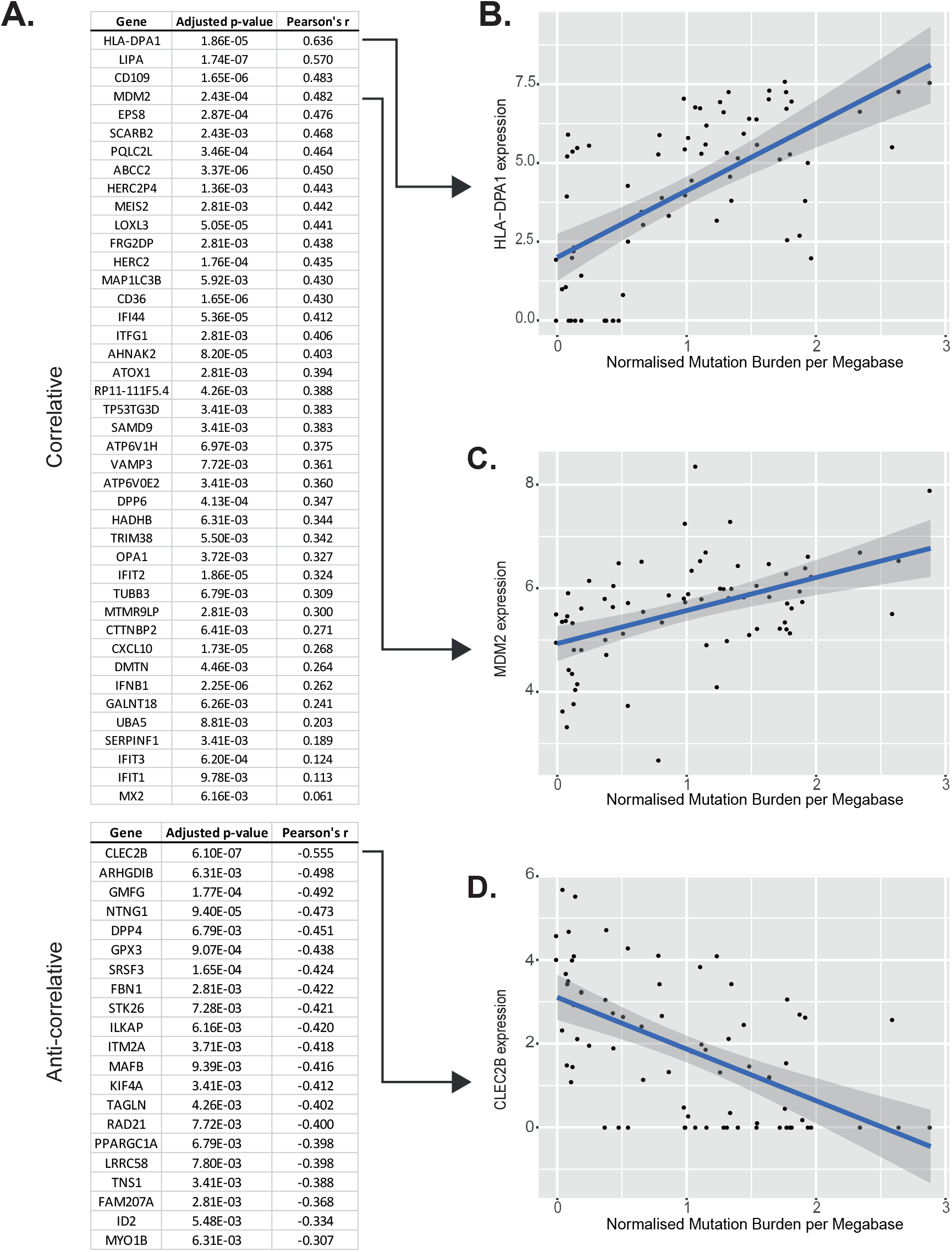
Differential expression analysis revealing genes significantly correlating with mutation burden. **A.** All genes differentially expressed from DeSeq2 analyses (see methods) with an adjusted p value < 0.01. The top correlative and anti-correlative genes are shown and ordered by the magnitude of their Pearson’s r value. **B-D.** Gene expression versus normalised mutation burden is shown for the top correlative and anti-correlative genes, as well as for *MDM2*, a gene of interest. Clones included in this analysis are from anatomic sites with greater than 3 standard deviations of mutation burdens among their cells, thus demonstrating a range of mutation burdens.

**Figure S4.**
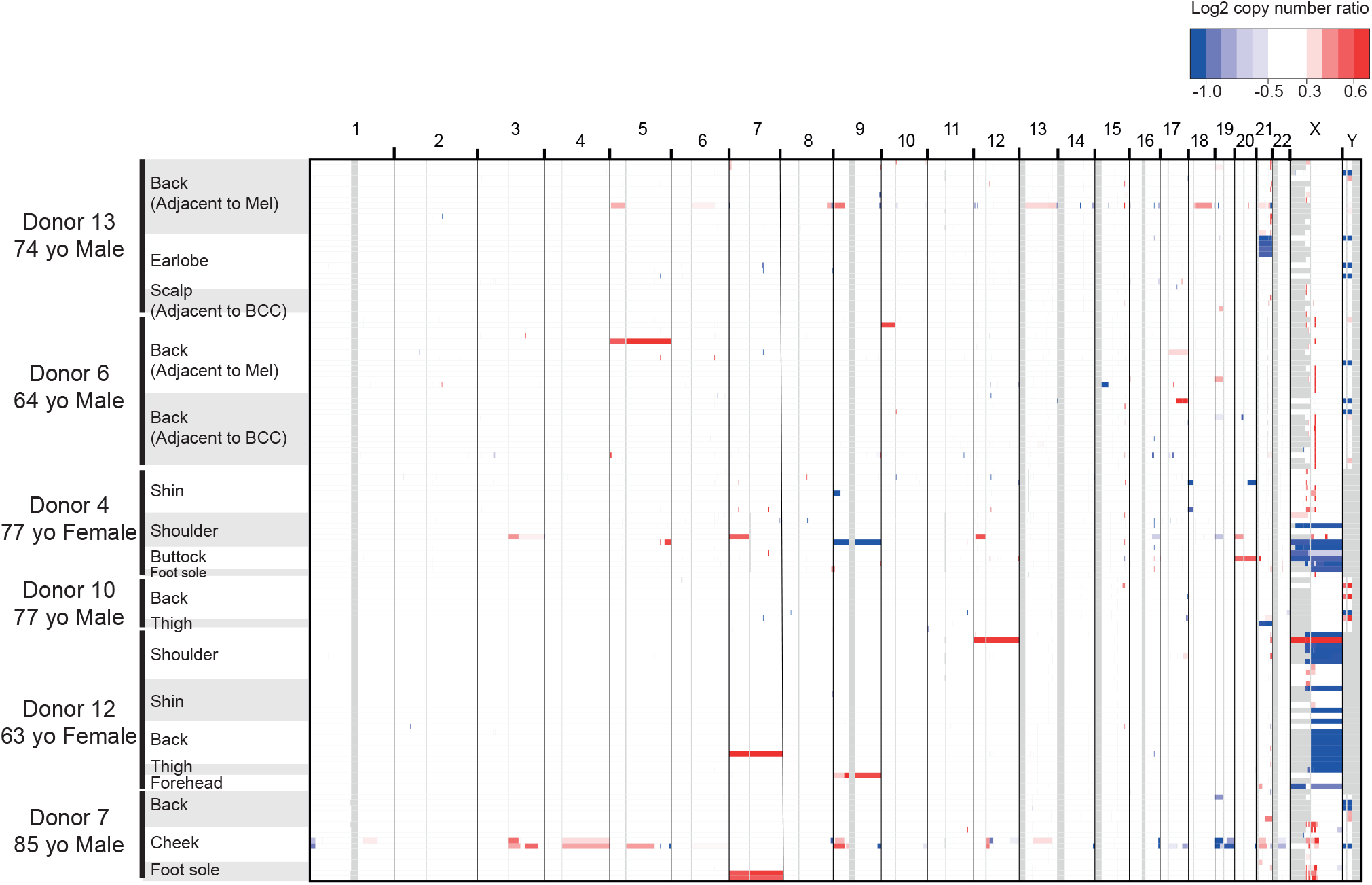
Copy number landscape of melanocytes from normal human skin. Copy number was inferred, as described, and segments (regions of equal copy number) are depicted, here, denoting gains (red) and losses (blue) for each melanocyte (rows). Note that copy number alterations over autosomes were rare, whilst the loss of one sex chromosome is a common occurrence. All X chromosome deletions in females affect the inactive X (see Table S4).

**Figure S5.**
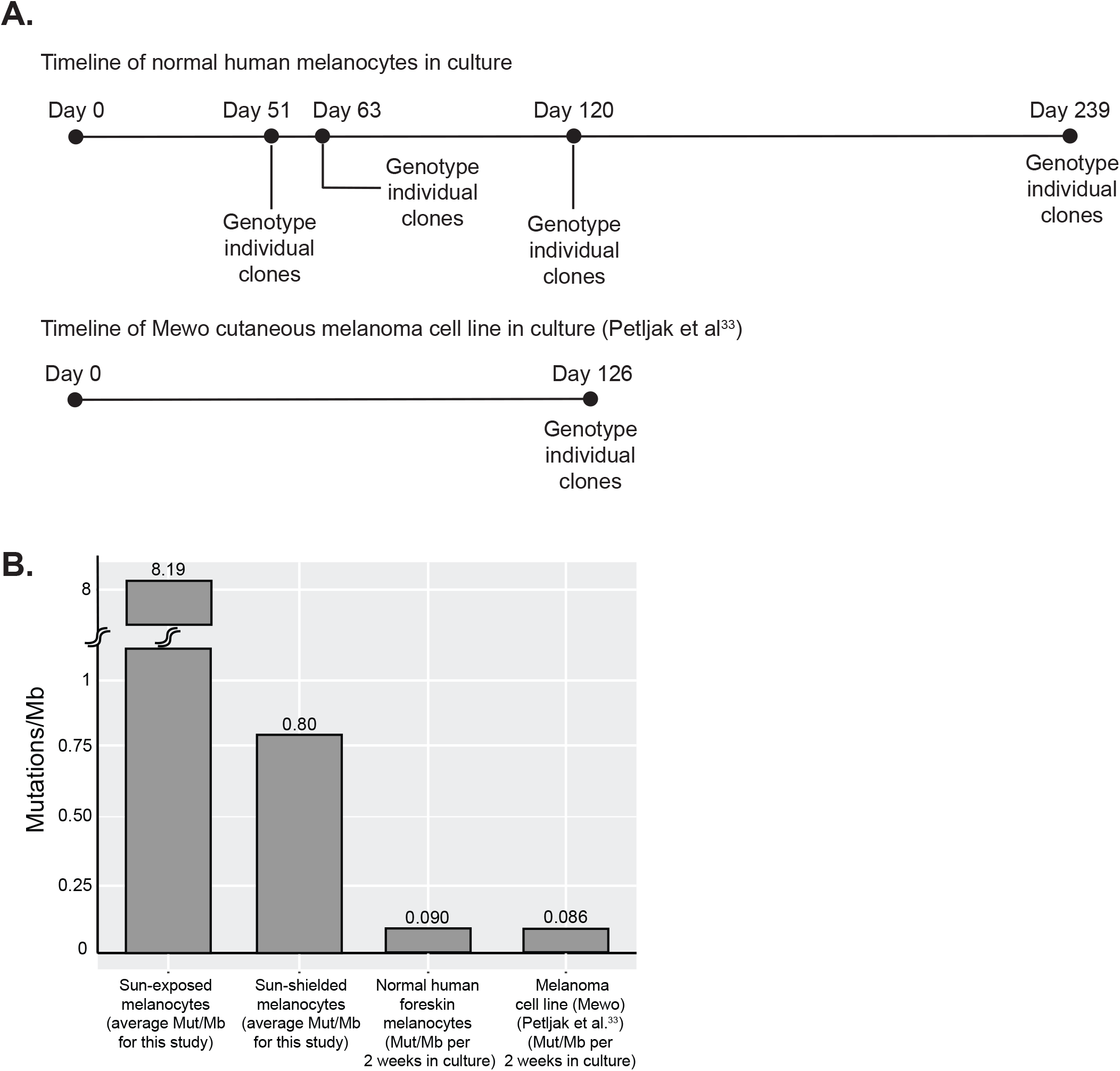
Melanocytes accumulate few mutations in tissue culture. **A.** We sequenced a bulk culture of neonatal melanocytes to establish the germline SNPs and somatic mutations in the dominant clones. We continued to passage the cell line for 239 days, genotyping individual clones at the timepoints indicated to establish the rate at which mutations were acquired in culture. In parallel, Petljak et al^33^ performed similar experiments on common cancer cell lines, and we analyzed their data from a melanoma cell line (Mewo) included in their study. **B.** On average, the mutation burden of neonatal melanocytes and Mewo cells respectively increased by 0.090 and 0.086 mutations/Mb (Mut/Mb) for every 2 weeks in tissue culture (we typically cultured melanocytes for 2 weeks in this study). To put these mutation burdens in perspective, the average mutation burdens of sun-exposed and sun-shielded melanocytes from this study are shown in comparison. Based on these results, we conclude that the brief period of tissue culture contributed little towards the mutation burdens observed in our study.

## Methods

### Skin tissue collection

Physiologically normal skin tissue was collected from cadavers (up to 8 days post-mortem) or from surgical discard tissue of living donors. Skin tissue from cadavers was collected from either the UCSF Autopsy program or the UCSF Willed Body Program. Family members consented to donate tissue from the UCSF Autopsy program, and Willed-Body donors consented to donate their tissues for scientific research prior to their expiration. Surgical discard tissue was collected from donors undergoing dermatologic surgery at UCSF, and their consent was obtained at the time of surgery. Donors from the UCSF Willed Body Program have consented to have any data derived from the donation to be deidentified, stored and shared securely, and used for research as required by the Federal Privacy Act of 1974, California Information Practices Act of 1977, and HIPAA (Health Insurance Portability and Accountability Act). Donors from clinical practice have consented to the release and sharing of deidentified clinical data and genetic testing information via HIPAA as guided by the NIH National Human Genome Research Institute.

Here, we define physiologically/clinically normal skin as skin lacking palpable or visible lesions. High resolution photos of each skin sample are available in the supplemental dataset. Skin tissue was stored at 4°C and processed in under 24h from time of collection.

### Establishment of epidermal skin cells in tissue culture

Skin tissue was briefly sterilized with 70% ethanol and rinsed with Hank’s Balanced Salt Solution (Thermo #14175095). Excess dermis was trimmed off and skin was cut into pieces (approximately 2×2 mm^2^) using surgical scalpel blades. Tissue was incubated in 10mg/ml dispase II (Thermo #17105-041) for 18hr at 4°C. The epidermis was peeled away from the dermis, incubated in 0.5% trypsin-EDTA (Thermo #15400-054) at 37°C for 4 min, and neutralized with 0.5mg/ml soybean trypsin inhibitor (Thermo #17075-029). Epidermal cells were plated in Medium 254 (Thermo #M254500) supplemented with human melanocyte growth supplement-2 (HGMS-2, Thermo #S0165) and antibiotic-antimycotic (Thermo #15240062). Cells were incubated at 37°C, 5% CO_2_ for 7-14 days.

### CRISPR engineering of a subset of cells

Initially, we presumed that it would be impossible to clonally expand single-cell sorted melanocytes from adult human skin, so we engineered mutations into the *CDKN2A* locus, as described^37^. This decision was based on our previous success in engineering *CDKN2A* mutations into foreskin melanocytes and our ability to clonally expand these melanocytes, thereby producing isogenetic population of engineered melanocytes^37^. However, during the course of these experiments, we recognized that control melanocytes, which were not engineered, could clonally expand under optimized tissue culture conditions, so we subsequently stopped engineering melanocytes. In total, 5 melanocytes were engineered prior to genotyping, as indicated in TableS1. Removal of these cell does not affect any of the conclusions from this study.

### Flow cytometry and cell culture of individual cell clones

Establishing epidermal cells in tissue culture produced a heterogeneous mixture of cells, comprised primarily of melanocytes and keratinocytes with some fibroblasts present. Differential trypsinization was used to separate melanocytes from keratinocytes using 0.05% trypsin-EDTA (Thermo #25300054) at 37°C for 2 min and 10 min, respectively. Trypsin was neutralized with 0.5mg/ml soybean trypsin inhibitor. Cells were centrifuged at 300 rpm for 5 min, resuspended in 300μl sorting buffer (1X PBS without Ca^2+^ and Mg^2+^ (Caisson Labs #PBL-01), 1mM EDTA (Thermo #AM9262), 25mM HEPES, pH 7.0 (Thermo #15630130), and 1% bovine serum albumin (Thermo #BP67110)), strained using test tube with 35μm cell strainer snap cap (Corning #352235), and single cell sorted into 96-well plates filled with 100μl complete Medium 254 using a Sony SH800S Cell Sorter. Cell sorting was performed using a 100μm microfluidic sorting chip with the 488nm excitation laser without fluorescent markers.

The next day, cells were screened to decipher their morphology and confirm that each well had only one cell. Individual melanocytes were grown in CnT-40 melanocyte medium (CELLnTEC #CnT-40) supplemented with antibiotic-antimycotic. A small number of cells had keratinocyte or fibroblast morphology. Keratinocytes were grown in 50:50 complete Medium 254 and Keratinocyte-SFM media (Thermo #17005042), and fibroblasts were grown in complete Medium 254 for 10-14 days. After 10-21 days, clone sizes ranged from 2-3000 cells (Table S1) and ceased any further expansion, prompting us to harvest these clones at their peak cell count. Approximately 37.5% of flow-sorted cells typically produced colonies, providing evidence that we are studying a prevalent and representative population.

### Extraction and amplification of DNA and RNA from each clone

Clones of 2-3000 cells do not yield enough genomic material to directly sequence using conventional library preparation technologies. For this reason, we elected to isolate both DNA and RNA from each clone and pre-amplify the nucleic acids prior to sequencing. To do this, we utilized the G&T-Seq protocol^14,15^.

G&T-Seq was performed, as described^14,15^. In brief, clones of cells were lysed in 7.5μl RLT Plus Buffer (Qiagen #1053393). mRNA and genomic DNA were separated using a biotinylated oligo d(T)_30_ VN mRNA capture primer (5′-biotin-triethyleneglycol-AAGCAGTGGTATCAACGCAGAGTACT30VN-3′, where V is either A, C or G, and N is any base; IDT) conjugated to Dynabeads MyOne Streptavidin C1 (Thermo #65001). cDNA was synthesized using the Smart-Seq2 protocol using SuperScript II reverse transcriptase (Thermo #18064014) and template-switching oligo (5′-AAGCAGTGGTATCAACGCAGAGTACrGrG+G-3′, where “r” indicates a ribonucleic acid base and “+” indicates a locked nucleic acid base; Qiagen). cDNA was amplified using KAPA HiFi HotStart ReadyMix kit (Roche #KK2502) and purified in a 1:1 volumetric ratio of Agencourt AMPure XP beads (Thermo #A63880). The average yield of amplified cDNA was 305ng. Genomic DNA was purified in a 0:0.72 volumetric ratio of Agencourt AMPure XP beads and amplified using multiple displacement amplification with the REPLI-g Single Cell Kit (Qiagen #150345) to yield an average of 815ng amplified genomic DNA per clone.

### Library preparation and next-generation sequencing of amplified DNA and RNA

We next prepared the amplified cDNA and amplified genomic DNA for sequencing. Library preparation was performed according to the Roche Nimblegen SeqCap EZ Library protocol. In brief, 250ng DNA input was sheared to 200bp using Covaris E220 in a Covaris microtube (Covaris #520077). End repair, A-tailing, adapter ligation (xGen Duel Index UMI adapters; IDT), and library amplification was performed using the KAPA HyperPrep kit (Roche #KK8504) and KAPA Pure Beads (Roche #KK8001). Library quantification was performed using the Qubit dsDNA High Sensitivity kit and quantitative PCR with the KAPA Quantification kit (Roche #KK4854) on a QuantStudio 5 real-time PCR system.

Target enrichment for next-generation sequencing was performed with the UCSF500 Cancer Gene Panel (developed by the UCSF Clinical Cancer Genomics Laboratory; Roche) or the SeqCap EZ Exome + UTR library probes (Roche #06740294001). All cells initially underwent targeted sequencing, and if a cell had a low mutation burden, or if a cell was phylogenetically related to other cells, we sequenced it again with exome baits. The exome sequencing data yielded more mutations, allowing us to infer mutational processes in low mutation burden cells and in distinct branches of phylogenetically related cells.

Hybridization reaction was performed using the SeqCap EZ Hybridization and Wash Kit (Roche #05634253001). xGen Universal blocking oligos (IDT #1075474), human COT 1 DNA (Thermo #15-279-011), and custom xGen Lockdown probes targeting the telomerase reverse transcriptase (*TERT*) promoter (IDT) were additionally added to the hybridization reaction. After library wash and PCR amplification, the captured library was quantified by Qubit and analyzed using the High Sensitivity DNA kit on Agilent’s Bioanalyzer 2500.

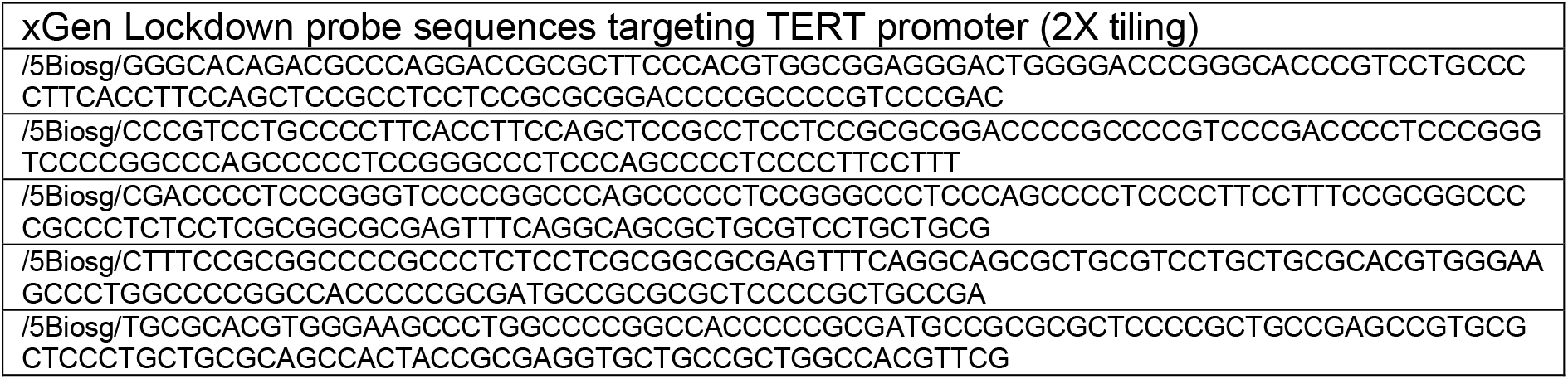
TERT promoter spike-in baits

Libraries were sequenced on an Illumina HiSeq 2500 or Novaseq (paired end 100bp or 150bp). On average, we achieved 489-fold unique coverage from targeted sequencing data, 86-fold unique coverage from exome sequencing data, and 7.75 million reads/clone from RNA-sequencing data.

### Data Deposition

Sequence data that support the findings of this study have been deposited in dbGaP (submission will be complete upon publication) with the accession code phs001979.v1.p1.

### Calling a preliminary set of variants

Variant call format files for each clone were generated as described^27,38^. Briefly, Fastq files underwent quality checks using FastQC and were subsequently aligned to the hg19 reference genome using the BWA-MEM algorithm (v0.7.13). BWA-aligned bam files were further groomed and deduplicated using Genome Analysis Toolkit and Picard. For each clone, variants were called using Mutect (v3.4.46) by comparing to bulk normal cells from a distant anatomic site. At this stage, the variants were composed primarily of amplification artifacts and somatic mutations. We leveraged the matched DNA/RNA sequencing data and haplotype information, detailed in the subsequent section, to distinguish between these entities.

### Harnessing the matched DNA/RNA sequencing data to remove amplification artifacts

The DNA and RNA from each clone were separately amplified, and consequently, amplification artifacts were unlikely to affect the same genomic coordinates in both the DNA- and RNA-sequencing reads (Fig. 1C). In contrast, somatic mutations should always overlap, assuming there was coverage of the mutant allele in both the DNA- and the RNA-sequencing data. We applied the following criteria to determine whether this assumption could be met.

To begin, we established rates of allelic dropout in our DNA- and RNA-sequencing data. From known heterozygous SNP sites, we empirically deduced that allelic dropout rates were less than 0.15% in our DNA-sequencing data. We achieved low levels of allelic dropout because of our high sequencing coverage, relatively uniform levels of coverage, and low levels of PCR-bias during amplification. Coverage in the RNA-sequencing data was more variable due to differences in gene expression, but from known heterozygous SNP sites, we empirically deduced that 15X coverage was sufficient to sample both alleles at nearly all variant sites. There were a small number of exceptions for which this did not hold true. Truncating mutations (nonsense, splice-site, and frameshift) are prone to nonsense-mediated decay and were commonly undersampled in our RNA-sequencing data relative to the wild-type allele. Also, mutations on the X-chromosome from female donors tended to be in 100% or 0% of RNA-sequencing reads, depending on whether they resided on the active or inactive X-chromosome. Aside from these examples, allelic variation in expression was minimal, particularly for highly expressed genes, as was previously reported^39^.

Based on these observations, a variant was considered a somatic mutation if it was present in both the DNA- and the RNA-sequencing data from the same clone. Conversely, a variant was considered an amplification artifact if the following conditions were met: the variant was present in the DNA-sequencing data but not the RNA-sequencing data, and there was at least 15X coverage in the RNA-sequencing data, and the variant was not truncating or on the X-chromosome. We declined to make a call in either direction for any variant that did not fulfil these conditions.

A limitation to this approach was that some variants did not reside in genes that were expressed. Nevertheless, 11.6% of variants could be classified as either a somatic mutation or amplification artifact by cross-validating the DNA/RNA sequencing data.

### Harnessing haplotype information to remove amplification artifacts

We also used haplotype information to distinguish between somatic mutations and amplification artifacts. Somatic mutations occur in *cis* with nearby germline polymorphisms, and this pattern is preserved during amplification (Fig. 1D). By contrast, amplification artifacts do not occur in complete linkage with nearby germline polymorphisms for the reasons described below (Fig. 1D).

The germline polymorphisms operate like unique molecular barcodes, designating which amplicons descended from each parental allele. The main reason why amplification artifacts are not in complete linkage with nearby polymorphisms is because there are multiple template molecules, associated with each parental allele, from which to amplify, and each template molecule can be amplified more than once – it is unlikely that the exact same mistakes are made during each independent amplification reaction over an error-free template. For example, we sequenced clonal expansions of cells, so each cell provided one molecule of double-stranded DNA from each allele. Furthermore, both strands of DNA are subject to amplification, thereby doubling the number of template molecules relative to the starting cell number. Finally, a single strand of DNA is repeatedly amplified during multiple displacement amplification, further enhancing the number of times an error-free template is utilized during amplification. Amplification artifacts therefore reveal themselves in the sequencing data by not occurring in complete linkage with nearby polymorphisms.

There was an exception for which the pattern described above did not hold true. A copy number gain or copy-number-neutral loss-of-heterozygosity (LOH) results in two or more copies of a single parental allele. If a somatic mutation occurs after the allelic duplication, then the somatic mutation would not be in complete linkage with nearby polymorphisms.

Consequently, we did not apply this methodology to root out amplification artifacts over regions of the genome for which there was an allelic duplication.

A limitation to this approach is that we used short-read sequencing technologies, so some variants were too far away from the nearest polymorphic sites to be phased. Nevertheless, 14.7% of variants could be classified as either a somatic mutation or amplification artifact, using the phasing approach.

### Inferring the mutational status of variants outside of the expressed or phase-able portions of the genome

In total, 25.1% of variants could be classified as either a somatic mutation or amplification artifact, using either the expression or the phasing approaches, described above. The remaining variants did not reside in portions of the genome that were sufficiently expressed or close enough to germline polymorphisms to permit phasing. For these variants, we inferred their mutational status from their variant allele frequency.

The majority of somatic mutations in our study were heterozygous, and these mutations, as expected, exhibited a normal distribution of mutant allele frequencies centered at 50% (Fig. 1E, S2B). The standard deviation of mutant allele frequencies in a given clone was dictated primarily by the number of starting cells, indicating that allelic biases, introduced during amplification, were the primary drivers of “noise” in our data.

By contrast, amplification artifacts exhibited a much different distribution of allele frequencies. Most amplification artifacts occurred in later rounds of amplification, and therefore had extremely low variant allele frequencies. However, a small number of amplification artifacts occurred in relatively early rounds of amplification and were disproportionately amplified thereafter. As a result, amplification artifacts exhibited a Poisson distribution of allele frequencies with a low peak but a long tail, sometimes extending into the range of allele frequencies seen for somatic mutations (Fig. 1E, S2B). As expected, the tail of this distribution was more extreme in clones with fewer starting cells because amplification biases were more exacerbated in these clones.

Due to the distinct distributions of variant allele frequencies for somatic mutations and amplification artifacts, a variant allele frequency cutoff could distinguish the vast majority of somatic mutations from amplification artifacts. However, the sensitivity and specificity of somatic mutation calls, using this approach, varied for each clone, primarily based on the clone size for the reasons described above. We were able to precisely define the sensitivity and specificity of mutation calls, and we could optimize the VAF cutoff for each clone by studying the overlap in variant allele frequencies from known somatic mutations and known amplification artifacts.

For each clone, we had a set of known somatic mutations and known amplification artifacts, situated in the expressed and phase-able portions of the genome. We were therefore able to determine the proportion of false positives and false negatives under the assumption that all variants above a given variant allele frequency were somatic mutations. Here, a “false positive” is an amplification artifact that would have been called somatic mutations, and a “false negative” is a somatic mutation that would have been called an amplification artifact. We plotted the sensitivity and specificity of mutation calls at different variant allele frequency cutoffs for each clone, and we chose the variant allele frequency cutoff that maximized these values – this value was then applied to the variants whose mutational status was unknown – i.e. the variants outside of the expressed and phase-able portions of the genome. For clones greater than 5 cells, we could typically infer somatic mutations at greater than 98% specificity and 98% sensitivity (Fig. S2C,D). We indicate in table S3 whether each mutation was validated or inferred by this approach.

### Copy Number Analysis

Copy number alterations were inferred from both the DNA- and the RNA-sequencing data using CNVkit^18,19^. We also integrated allelic frequencies from somatic mutations and germline heterozygous SNPs.

First, we inferred copy number alterations from the DNA-sequencing data. CNVkit can be run in reference or reference-free mode. We elected to run CNVkit in reference mode, and in doing so, we created several references, encompassing panels of clones without copy number alterations that were amplified and prepared for sequencing in similar batches. This approach consistently produced the least noisy copy number profiles, as compared to reference-free mode or a universal reference. All other parameters were run on their default settings.

Second, we inferred copy number alterations from the RNA-sequencing data. Briefly, CNVkit assumes the expression of a gene correlates with its copy number status. Of course, the expression of a gene is dictated by several factors, including, but not limited, to copy number. As an input, CNVkit accepts correlation values from an independent dataset between expression and copy number. Here, we included correlation values from the melanoma TCGA project. Given this input, CNVkit downweighs genes whose expression does not correlate well with copy number.

Third, we calculated allelic imbalance over germline heterozygous germline SNPs. Copy number alterations are expected to induce imbalances over these sites. Additionally, we calculated the allelic frequencies of somatic mutations across the genome, as these, too, would be modulated by copy number alterations.

Finally, we manually reviewed the copy number and variant allele information to call copy number alterations that were supported by each approach.

### An overview of the genetic landscape of each sequenced melanocyte

We have included individual summaries of the 133 sequenced clones which describes cellular morphology and tissue images, validation of variant allele fraction of raw calls, copy number alterations, and CDKN2A status (where applicable) (https://figshare.com/s/bb5614d5ab4554516278). Scripts and resources to perform analyses downstream of variant calling are available on GitHub (https://github.com/elliefewings/Melanocytes_Tang2020).

### Admixture Analysis (related to Fig. S1A)

Bulk normal cells were analyzed to identify germline variants present in each studied donor. Donor ethnicity was inferred via Admixture analysis using a Bayesian modelling approach employed by the tool STRUCTURE (v2.3.4)^40^. A set of 7662 common variants (1000 genomes population allele frequency > 0.05) with a sequencing depth of greater than 10 across all donors and all 2504 samples from the 1000 genomes study^41^ were selected. The burn-in period and analysis period were both completed with 10,000 repetitions as per the tool recommendations to achieve accurate estimations of admixture. To select an appropriate number of populations (*K)*, the algorithm was run using *K* estimations of 5 to 9. A final *K* value of 8 was selected to appropriately cluster populations without overfitting. The data were plotted using the STRUCTURE GUI plotting tool. Ethnicity of donors within this study was inferred by their similarity to known populations within the 1000 genomes set^41^.

### RNA gene expression analysis (related to figure 1B and S1B)

RNA sequencing reads were aligned to the transcriptome as well as the hg19 reference genome using STAR alignment tool (v2.5.1b)^42^. Transcripts were quantified using RNA-Seq by Expectation-Maximization (RSEM)^43^ and filtered to remove those with fewer than 10 reads across all clones as recommended by DESeq2 R package documentation. A variance stabilizing transformation was applied to the data and a Barnes-Hut T-distributed stochastic neighbour embedding (t-SNE) algorithm was performed to cluster related cells on the expression of the top 500 genes using the Rtsne R package (v0.15) with a perplexity of 6 over 1000 iterations.

Differential expression analysis was completed on the quantified transcript values using DESeq2 R package^44^ (v1.22.2). Three experimental designs were produced, selecting for differentially expressed genes that are over-expressed in fibroblasts, melanocytes, and keratinocytes independently. The data were log_2_ transformed and a heatmap was generated presenting the top 20 significantly differentially over-expressed genes per cell type.

Gene set enrichment analysis was performed across the significantly differentially over-expressed genes from each cell type using the Molecular Signatures Database (v6.2) webtool. The top significantly enriched pathways were examined for their relation to the cell-type of interest.

### Mutation burden and signature analysis (related to figure 2)

The mutation burdens reported in figure 2 correspond to the number of somatic mutations in a given clone divided by the genomic footprint for which mutations could be detected. Due to differences in depth of coverage across bam files and the unevenness of coverage in a given bam file, mutations were not callable at every base present in target region. Additionally, we used both a targeted and exome sequencing panel in this study, which produce two different sequencing footprint sizes. To account for these issues, we calculated callable sequencing footprints for each clone and corresponding reference. On-target bam files were created per clone and per bulk normal. The coverage of each on-target base was calculated using the bedtools (v2.25.0) genomecov command, and the number of bases covered by more than 5 reads was counted in each bam file. The minimum value between a clone’s bam and its reference bam was used as the footprint from which to calculate a mutation burden for each clone.

To perform mutational signature analysis, surrounding genomic contexts were applied to single nucleotide variants identified in each clone using the Biostrings hg19 human genome sequence package (BSgenome.Hsapiens.UCSC.hg19 v1.4.0). Variant contexts were used to assess the proportion of each clone’s mutational landscape that could be attributed to a mutagenic process using the deconstructSigs R package (v1.8.0). A set of 48 signatures recently described by Petljak *et al*^33^ were analysed, with particular attention paid to the single base substitution signatures 7a, 7b, and 7c that are associated with ultraviolet light exposure.

### Gene expression correlation with mutation burden (related to figure S3)

RNA data was used to explore the variability in mutation burdens, often observed over a single site. Sites with greater than 3 standard deviations of mutation burdens, demonstrating the presence of both high and low mutation burden clones, were selected for analysis. Mutation burdens were normalized to the median of each anatomic site. Differential expression analysis was then performed using DESeq2 R package^44^ (v1.22.2). Genes with expression changes significantly associated (adjusted p value < 0.01) with a continuous change in mutation burden are highlighted in figure S3.

### Estimating mutation acquisition over time in tissue culture (related to “Melanocytes can persist as fields of related cells within the skin” section of the results)

We established skin cells in tissue culture for 7-14 days prior to single-cell sorting and clonal expansion. Any mutation that arose after clonal expansion would be recognizable since it would only be present in a proportion of daughter cells, thus appearing subclonal. However, mutations that arose during the brief period of tissue culture preceding clonal expansion could be mistaken as a mutation that occurred while the cell was still situated in the skin. We therefore sought to establish the rate at which melanocytes accumulate *de novo* mutations in tissue culture to determine whether this was a meaningful contribution to the total mutation burden that we observed in our cells.

Towards this goal, we followed the framework recently put forth by Petljak and colleagues^33^ – in that study, the authors sequenced subclones of daughter cells from common cancer lines at different generational time points for up to 161 days, thereby revealing the mutational processes operating during their time in tissue culture. Here, we sequenced a bulk culture of normal human melanocytes derived from human foreskin to establish the germline variants and somatic mutations in the dominant clones. We continued to culture these cells, and at time points of 51, 63, 120, and 239 days, we single-cell sorted and clonally expanded individual cells. We genotyped each clonal expansion, following the same protocol that was applied to melanocytes in this study. From these analyses, we estimate that mutations occur at a rate of .045 mutations/Mb per 7 days in tissue culture. To put this in perspective, the mutation burden of melanocytes from the bottom of the foot was .25 mutations/Mb. Based on these findings, we conclude that the number of mutations accumulated in tissue culture was negligent as compared to the number of mutations that pre-existed in melanocytes that were profiled for this study.

We also analyzed the publicly available data from Petljak *et al*^33^ to deduce the rate at which melanoma cell lines accumulate mutations in tissue culture. From these analyses we estimate that mutations occur at a rate of .043 mutations/Mb per 7 days – in line with our estimates for normal human melanocytes.

Taken together, it is not surprising that the number of mutations collected from 7 days in tissue culture is negligible as compared to the number of mutations collected from decades situated in the skin.

### Phylogenetic tree construction (related to figure 4)

Pairwise comparisons of melanocyte mutation calls were performed to identify sets of melanocytes with shared mutations, and when this occurred, phylogenetic trees were constructed from the shared and unshared mutations. In figure 4, trunk lengths correspond to the number of shared mutations, and branch lengths correspond to the number of unshared mutations. If there was an allelic deletion in one clone, we did not assign mutations in the clone lacking the deletion over the deletion area to the branch. Shared mutations were discarded if there was insufficient coverage in the reference to rule out the possibility that the mutation was a germline SNP. Unshared mutations were discarded if there were trace reads in the clone for which the mutation was not detected or if sequencing coverage was insufficient in both clones to definitively make a call. In practice, few mutations needed to be discarded by these filtering criteria because we achieved high sequencing coverage in our clones.

